# Assessing Genetic Diversity in Critically Endangered *Chieniodendron hainanense* Populations within Fragmented Habitats in Hainan

**DOI:** 10.1101/2023.04.24.538051

**Authors:** Li Zhang, Hai-Li Zhang, Yukai Chen, Mir Muhammad Nizamani, Tingtian Wu, Tingting Liu, Qin Zhou

## Abstract

Habitat fragmentation engenders a reduction in the geographic distribution of species, thereby rendering diminutive populations susceptible to extinction due to environmental, demographic, and genetic factors. *Chieniodendron hainanense* (henceforth *C. hainanense*) exemplifies a wild plant with extremely small populations (WPESP) and faces endangerment, necessitating urgent national conservation efforts. Elucidating the genetic diversity of *C. hainanense* is crucial for uncovering underlying mechanisms and devising protective strategies. In the present study, 35 specimens from six distinct cohort groups were genotyped utilizing genotyping-by-sequencing (GBS) and single nucleotide polymorphism (SNP) methodologies. The results indicated that *C. hainanense* exhibits limited genetic diversity. Observed heterozygosity within *C. hainanense* populations spanned from 10.79% to 14.55%, with an average value of 13.15%. The six *C. hainanense* populations can be categorized into two distinct groups: (1) Diaoluoshan and Baishaling, and (2) Wuzhishan, Huishan, Bawangling, and Jianfengling. The degree of genetic differentiation among *C. hainanense* populations is relatively weak. The observed loss of diversity can be attributed to the effects of natural selection.

## 1 Introduction

The conservation of biodiversity is crucial for sustaining the stability and resilience of ecosystems. Genetic diversity constitutes a pivotal element in the long- term persistence of species, particularly for those facing extinction risks. *Chieniodendron hainanense* represents a rare and imperiled plant species characterized by extremely small populations (WPESP) and is endemic to the fragmented habitats of Hainan, China. Owing to the adverse consequences of habitat fragmentation, the genetic diversity of *C. hainanense* populations warrants significant attention in conservation initiatives. Comprehending the genetic architecture and diversity of these populations enables the development of suitable management approaches and facilitates species restoration efforts.

The genetic structure and diversity of a species play critical roles in their ability to withstand adverse environments and evolve over time. Studying the genetic makeup of wild plants with extremely small populations can provide insights into their evolutionary history, population dynamics, and response to environmental changes, enabling the development of effective conservation strategies (Yang et al., 2018; Zhang et al., 2021). Such research is also essential for advancing DNA sequencing technology and conservation biology (Windig et al., 2010).

Habitat fragmentation and overuse of resources have greatly impacted species’ genetic diversity, survival, adaptability, and biodiversity (Su et al., 2020). Habitat fragmentation reduces genetic diversity by isolating populations and restricting gene exchange, leading to an increase in genetic drift and inbreeding depression (Bijlsma et al., 2012; Wang et al., 2022). The reduced genetic diversity weakens the ability of species to adapt to environmental changes and increases the risk of species extinction (Luquet et al., 2012). Habitat fragmentation also reduces the number of plant populations, increases spatial isolation, and hinders the maintenance of population genetic diversity by disrupting dispersal, gene exchange, and inter-species interactions (Aguilar et al., 2008).

Wild plants with extremely small populations (WPESP) are endangered species under national key protection and require urgent rescue efforts. Their population sizes are smaller than the minimum viable and are usually distributed in a narrow area, making them highly vulnerable to extinction (Zang, 2020; Jain et al., 2013). The Chinese government has launched the National Project Plan for the Rescue and Protection of Wild Plants in Extremely Small Populations (2011-2015) to protect 120 WPESP species, most of which are endemic to China and have significant ecological, scientific, cultural, and economic value (Ma et al., 2013; Yang et al., 2020). The loss of biological and genetic values of WPESP due to their extinction can have significant adverse effects on human society and the ecosystem (Zhang et al., 2018). Therefore, it is crucial to prioritize research on WPESP conservation in current biodiversity conservation studies in China.

Chieniodendron hainanense is a second-class national key protected wild plant and a unique member of the Annonaceae family Chieniodendron genus in China. The species is an evergreen tree that grows up to 16 m tall with a DBH of about 50 cm, and its distribution is limited to the Guangxi Zhuang Autonomous Region and some areas of Hainan Province. Habitat fragmentation due to human activities such as logging has drastically reduced the distribution area of C. hainanense, and the wild resources of its population are scarce. Field surveys have shown that the existing populations of C. hainanense are mainly distributed in nine primary forest areas dominated by fragmented secondary rainforests in Hainan, with many original populations already disappeared (Jiang et al., 2021). The species has poor self-renewal ability and is highly sensitive to external disturbances, making its wild resources already endangered. Although some research reports have focused on the population structure, dynamic characteristics, leaf morphology, petal nodules development, and functional biochemical activities of C. hainanense, very few studies have investigated its endangerment mechanisms at the molecular level. To develop targeted and comprehensive conservation strategies, it is necessary to understand the causes of its endangerment, including its genetic diversity, and related components (Sork et al., 2006).

In this investigation, our objective is to evaluate the genetic diversity and population structure of C. hainanense across its fragmented habitats in Hainan, utilizing genotyping-by-sequencing (GBS) and single nucleotide polymorphism (SNP) techniques. Through the analysis of 35 specimens from six discrete cohort groups, we will scrutinize patterns of genetic variation, differentiation, and inbreeding both within and among the populations. The outcomes of this research will yield valuable insights into the conservation status and management of C. hainanense, promoting the formulation of efficacious strategies for the preservation and restoration of this critically endangered species. Additionally, our findings will augment the understanding of the wider implications of habitat fragmentation on genetic diversity in plant populations and inform conservation endeavors for other threatened plant species inhabiting fragmented ecosystems.

## 2 **Materials and methods**

### 2.1 Study area

Hainan Island (E108°37′-111°03′, N18°10′-20°10) is situated in the southern part of China, on the northern edge of the tropics (Figure 1). It has a tropical monsoon climate and a mild climate with long summers and short winters (Zhang et al., 2022). The annual average temperature ranges from 22°C to 27°C, and the island has abundant water. Hainan Island is an important distribution area for China’s monsoon forests and tropical rainforests (Zhang et al., 2022). The main forest vegetation types in Hainan Island are tropical rainforest, evergreen broad-leaved forest, coniferous forest, and plantation forest. Hainan Province has the largest tropical rainforest in China, with rich flora and fauna resources, covering 42.5% of the total tropical area of the country (Li et al., 2022). The critical areas for biodiversity conservation in my country, “National Of the 120 species of wild plants in extremely small populations identified in the Plan for the Rescue and Protection of WPESP (2011- 2015). Hainan has 24 key protection targets. Yunnan and Hainan are critical areas for the protection of WPESP (Sun et al., 2022)

**Figure 1.**
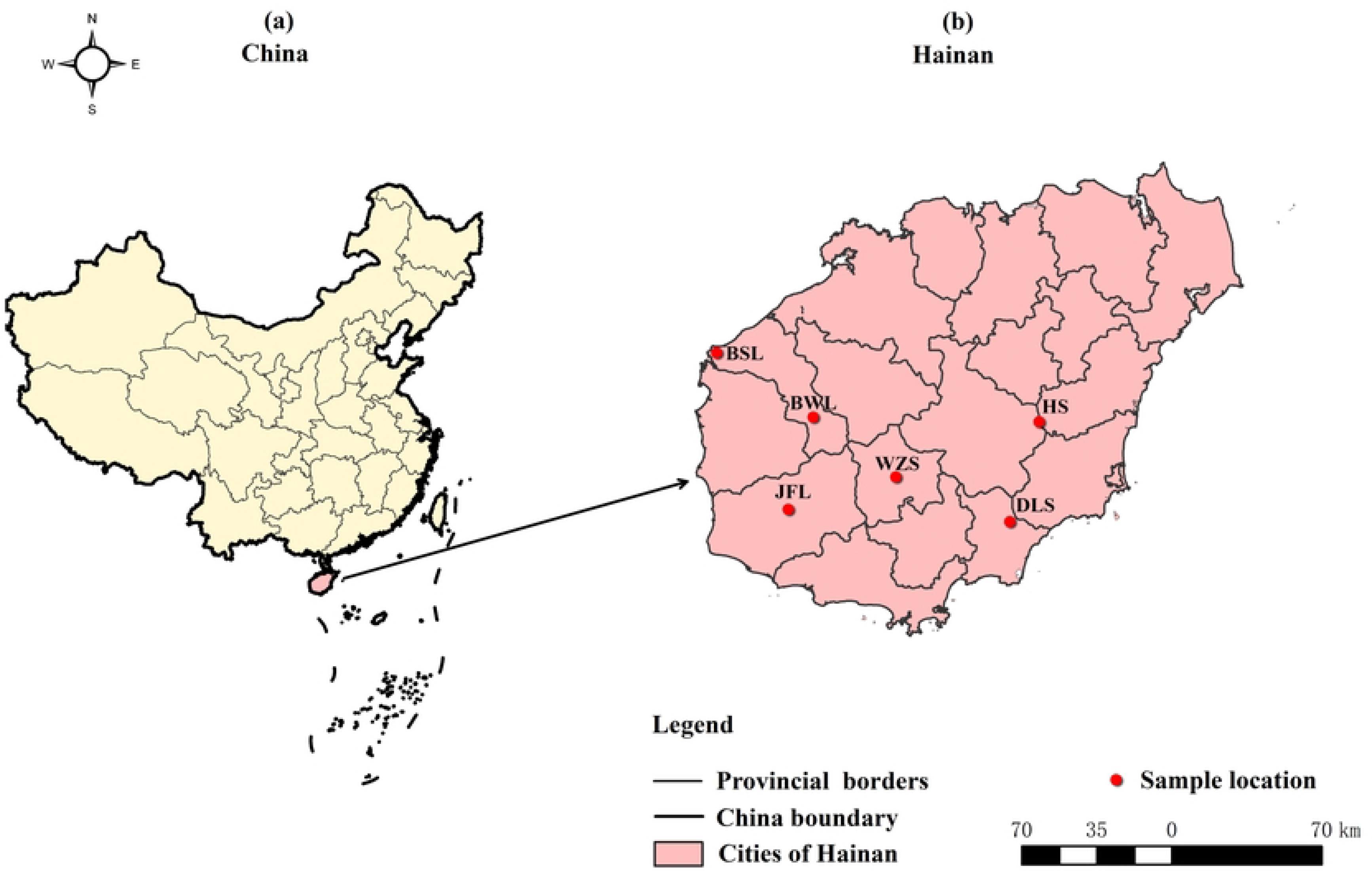
Distribution of *C. hainanense* on Hainan Island and sample collection information map. (a) Location of Hainan Island in China, (b) Distribution of *C. hainanense* on Hainan Island.

### 2.2. **Materials**

During 2019-2021, 35 *C.hainanen* samples were continuously collected from Hainan Island. They were divided into six regions according to the geographical origin of the samples, namely, the Bawangling population (samples BWL_1-BWL_8), the Jianfengling population (JFL_1-JFL_5), the Luoshan population (DLS_1-DLS_10), Huishan population (HS_1-HS_6), Wuzhishan population (WZS_1-WZS_3), Baishaling population (BSL_1-BSL_3). The division of regional populations, latitude, longitude, and the number of samples are shown in Table 1.

**Table 1.** The list of information for *C. hainanense* sample collection on Hainan Island

| Area code | Location | Longitude | Latitude | Sample | Number of samples |
| --- | --- | --- | --- | --- | --- |
| BWL | Bawanglin | 109°07'-109°12'E | 19°04'-19°07'N | BWL_1-BWL_8 | 8 |
| JFL | Jianfengling | 109°39'-109°48'E | 18°41'-18°46'N | JFL_1-JFL_5 | 5 |
| DLS | Diaoluoshan | 109°19'-109°56'E | 19°03'-19°04'N | DLS_1-DLS_10 | 10 |
| HS | Huihan | 110°08'-110°09'E | 19°03'-19°04'N | HS_1-HS_6 | 6 |
| WZS | Wuzhishan | 108°50' | 18°46'N | WZS_1-WZS_3 | 3 |
| BSL | Baishalin | 109°14'E | 19°11'N | BSL_1-BSL_3 | 3 |

### 2.3 Methods

Firstly, DNA was extracted from 35 *C. hainanense* leaf samples, and the quality and concentration of the extracted DNA were tested before being sent to Hangzhou Lianchuan Biotechnology Co. After sequencing was completed, the SNPs of the *C. hainanense* genome were mined. Based on these SNPs, the phylogenetic tree analysis of *C. hainanense* was obtained using the neighbor-joining algorithm of MEGA software. Principal component analysis (PCA) was then performed on *C. hainanense* populations based on the SNPs. Additionally, the population structure of all samples was analyzed using admixture software to obtain the distribution of genetic material in different populations of *C. hainanense*. Finally, genetic distances among all samples were calculated based on the SNPs. The detailed method is described in the paper by Chen et al. (2022).

#### 2.3.1 **Enzyme digestion protocol design**

Our simplified genome digestion scheme selected according to other research methods is as follows, the restriction enzyme combination of HaeIII + Hpy166II was selected. The ’Insert Size ’ was selected as ’550-600bp’ (Xia et al., 2019).

#### 2.3.2 **Sequencing Quality Control**

The Raw data (the number of reads in the original downstream data) generated by sequencing is pre-processed by quality filtering to obtain CleanData. The specific processing steps are as follows: 1) remove the adapter, 2) remove the reads containing N (N means the information of bases cannot be determined) with a proportion of more than 5%, 3) remove the low-quality reads (the number of bases with quality value Q<=10 accounts for more than 20% of the whole reads), 4) count the raw sequencing volume, effective sequencing volume, Q20 (the proportion of bases with quality values greater than or equal to 20, sequencing error rate less than 0.01), Q30 (the proportion of bases with quality values greater than or equal to 30, sequencing error rate less than 0.001), GC means guanine (G) and cytosine (C)content, and perform a comprehensive evaluation.

#### 2.3.3 **Comparison of consistency sequences**

We used Burrows-Wheeler aligner (BWA) software to match the sequencing data to the consistent sequences obtained from reads clustering. Since the reference used is the consistency sequence obtained from reads clustering, the matching rate will vary somewhat between samples.

#### 2.3.4 Variation detection and SNP statistics

After comparing the data with the concordant sequences, we used Genome Analysis Toolkit (GATK) and SAMtools software for variant detection, retaining the SNPs that were consistently output by both software as reliable loci. We further processed the SNP data by filtering them based on MAF > 0.05 and data integrity > 0.8 and retained the SNPs with polymorphisms among them. The final filtered SNPs were input to the subsequent evolutionary analysis.Based on the obtained SNP data, we analyzed the genetic evolution and structure of the population using the differences in genetic information among the samples of *C. hainanense*, including the phylogenetic relationships among the samples, population structure, principal component analysis (PCA), and relatedness among the samples. The following part of the analysis involves grouping samples, and the 35 samples were divided into six groups according to species for analysis.

## 3 **Results**

### 3.1 Simplified genome quality inspection of *C. hainanense*

A total of 49.16 GB of raw data were obtained through genotyping-by-sequencing (GBS) on 35 *C. hainanense* samples (Table 2). After removing the base information that could not be determined in the adapter sequence, the remaining 48.52 GB of high-quality sequencing data (Clean data), the average of each sample was 1.39 GB. In the simplified genome of *C. hainanense,* the average Q20 was 96.71%; the average Q30 was 90.97%. The average ratio (GC content) of guanine (G) and cytosine (C) is 40.60%. The results indicated that the sequencing quality of Q20 and Q30 was high, the GC distribution was reasonable, and the sequencing information was reliable.

**Table 2.**
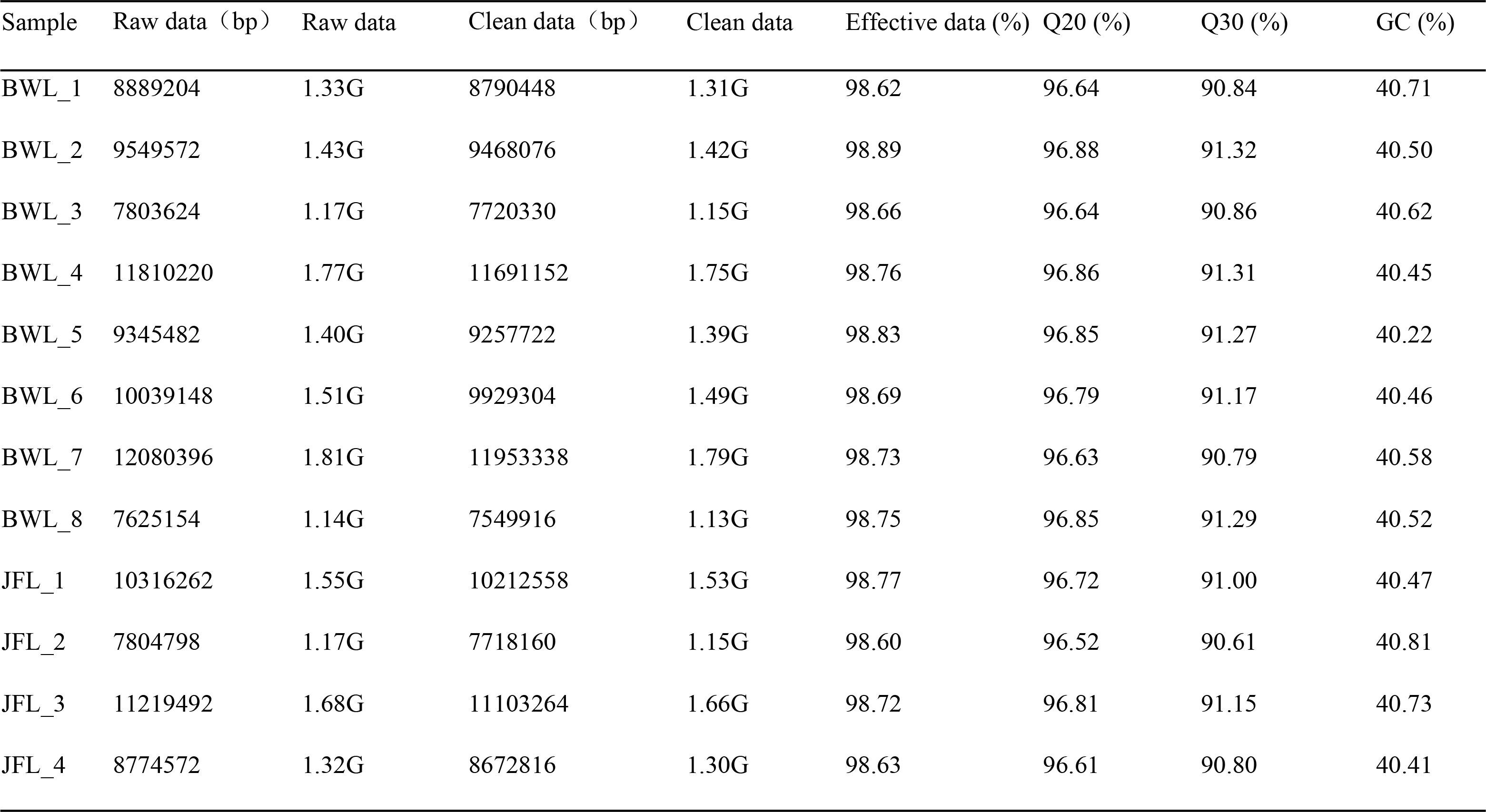
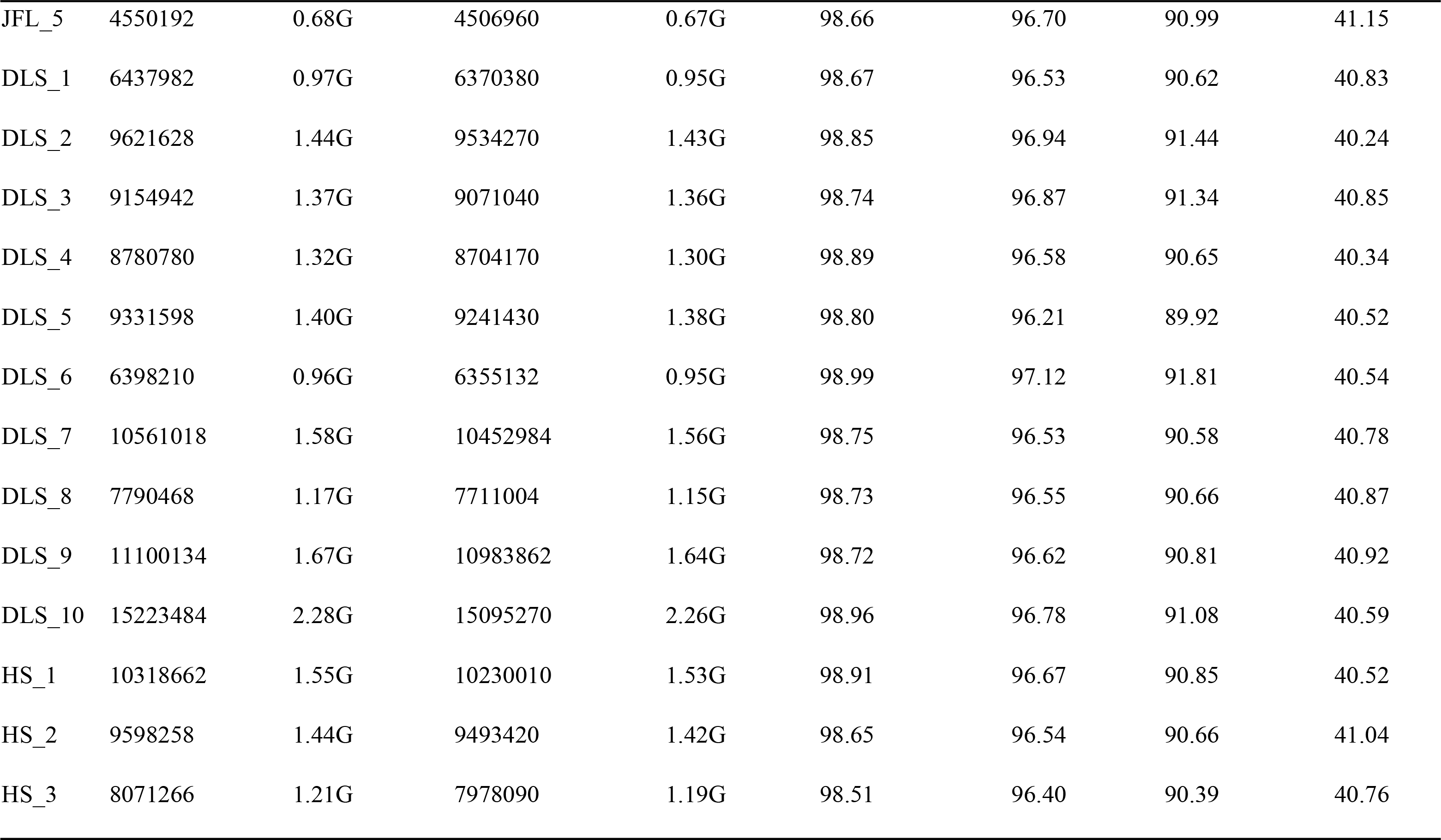
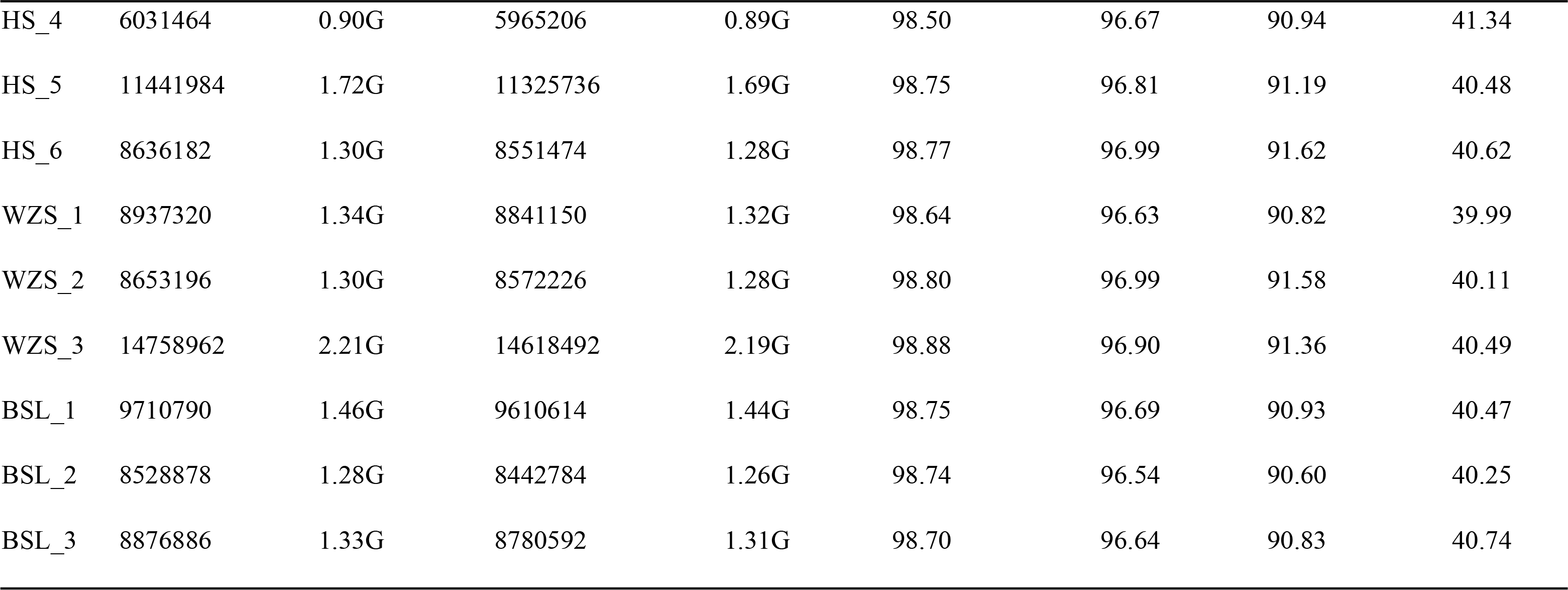
Description of the materials used and the GBS dataset

| Sample | Raw data (bp) | Raw data | Clean data (bp) | Clean data | Effective data (%) | Q20 (%) | Q30 (%) | GC (%) |
| --- | --- | --- | --- | --- | --- | --- | --- | --- |
| BWL_1 | 8889204 | 1.33G | 8790448 | 1.31G | 98.62 | 96.64 | 90.84 | 40.71 |
| BWL_2 | 9549572 | 1.43G | 9468076 | 1.42G | 98.89 | 96.88 | 91.32 | 40.50 |
| BWL_3 | 7803624 | 1.17G | 7720330 | 1.15G | 98.66 | 96.64 | 90.86 | 40.62 |
| BWL_4 | 11810220 | 1.77G | 11691152 | 1.75G | 98.76 | 96.86 | 91.31 | 40.45 |
| BWL_5 | 9345482 | 1.40G | 9257722 | 1.39G | 98.83 | 96.85 | 91.27 | 40.22 |
| BWL_6 | 10039148 | 1.51G | 9929304 | 1.49G | 98.69 | 96.79 | 91.17 | 40.46 |
| BWL_7 | 12080396 | 1.81G | 11953338 | 1.79G | 98.73 | 96.63 | 90.79 | 40.58 |
| BWL_8 | 7625154 | 1.14G | 7549916 | 1.13G | 98.75 | 96.85 | 91.29 | 40.52 |
| JFL_1 | 10316262 | 1.55G | 10212558 | 1.53G | 98.77 | 96.72 | 91.00 | 40.47 |
| JFL_2 | 7804798 | 1.17G | 7718160 | 1.15G | 98.60 | 96.52 | 90.61 | 40.81 |
| JFL_3 | 11219492 | 1.68G | 11103264 | 1.66G | 98.72 | 96.81 | 91.15 | 40.73 |
| JFL_4 | 8774572 | 1.32G | 8672816 | 1.30G | 98.63 | 96.61 | 90.80 | 40.41 |
| JFL_5 | 4550192 | 0.68G | 4506960 | 0.67G | 98.66 | 96.70 | 90.99 | 41.15 |
| DLS_1 | 6437982 | 0.97G | 6370380 | 0.95G | 98.67 | 96.53 | 90.62 | 40.83 |
| DLS_2 | 9621628 | 1.44G | 9534270 | 1.43G | 98.85 | 96.94 | 91.44 | 40.24 |
| DLS_3 | 9154942 | 1.37G | 9071040 | 1.36G | 98.74 | 96.87 | 91.34 | 40.85 |
| DLS_4 | 8780780 | 1.32G | 8704170 | 1.30G | 98.89 | 96.58 | 90.65 | 40.34 |
| DLS_5 | 9331598 | 1.40G | 9241430 | 1.38G | 98.80 | 96.21 | 89.92 | 40.52 |
| DLS_6 | 6398210 | 0.96G | 6355132 | 0.95G | 98.99 | 97.12 | 91.81 | 40.54 |
| DLS_7 | 10561018 | 1.58G | 10452984 | 1.56G | 98.75 | 96.53 | 90.58 | 40.78 |
| DLS_8 | 7790468 | 1.17G | 7711004 | 1.15G | 98.73 | 96.55 | 90.66 | 40.87 |
| DLS_9 | 11100134 | 1.67G | 10983862 | 1.64G | 98.72 | 96.62 | 90.81 | 40.92 |
| DLS_10 | 15223484 | 2.28G | 15095270 | 2.26G | 98.96 | 96.78 | 91.08 | 40.59 |
| HS_1 | 10318662 | 1.55G | 10230010 | 1.53G | 98.91 | 96.67 | 90.85 | 40.52 |
| HS_2 | 9598258 | 1.44G | 9493420 | 1.42G | 98.65 | 96.54 | 90.66 | 41.04 |
| HS_3 | 8071266 | 1.21G | 7978090 | 1.19G | 98.51 | 96.40 | 90.39 | 40.76 |
| HS_4 | 6031464 | 0.90G | 5965206 | 0.89G | 98.50 | 96.67 | 90.94 | 41.34 |
| HS_5 | 11441984 | 1.72G | 11325736 | 1.69G | 98.75 | 96.81 | 91.19 | 40.48 |
| HS_6 | 8636182 | 1.30G | 8551474 | 1.28G | 98.77 | 96.99 | 91.62 | 40.62 |
| WZS_1 | 8937320 | 1.34G | 8841150 | 1.32G | 98.64 | 96.63 | 90.82 | 39.99 |
| WZS_2 | 8653196 | 1.30G | 8572226 | 1.28G | 98.80 | 96.99 | 91.58 | 40.11 |
| WZS_3 | 14758962 | 2.21G | 14618492 | 2.19G | 98.88 | 96.90 | 91.36 | 40.49 |
| BSL_1 | 9710790 | 1.46G | 9610614 | 1.44G | 98.75 | 96.69 | 90.93 | 40.47 |
| BSL_2 | 8528878 | 1.28G | 8442784 | 1.26G | 98.74 | 96.54 | 90.60 | 40.25 |
| BSL_3 | 8876886 | 1.33G | 8780592 | 1.31G | 98.70 | 96.64 | 90.83 | 40.74 |

### 3.2 *C. hainanense* SNP site mining

In this study, the heterozygosity of the six populations of *C. hainanense* ranged from 10.79% to 14.55%, and the average heterozygosity is 13.15%, which shows the genetic diversity of *C. hainanense* trees is low (Table 3).

**Table 3.** SNP Statistical results

| Population code | SNP number | Heter LociNum | Homo LociNum | Hetloci-ratio |
| --- | --- | --- | --- | --- |
| BWL | 181002 | 25265 | 155737 | 13.93% |
| JFL | 171854 | 23417 | 148437 | 13.45% |
| DLS | 183868 | 25185 | 158683 | 13.59% |
| HS | 199196 | 25077 | 174119 | 12.56% |
| WZS | 190463 | 27590 | 162873 | 14.55% |
| BSL | 172311 | 18612 | 153699 | 10.79% |

### 3.3 Genetic evolution and population analysis

#### 3.3.1 Phylogenetic evolutionary tree

We divided the 35 *C. hainanense* samples roughly into two general taxa, which can be further subdivided into several subgroups. Group I mainly comprise resources from Diaoluoshan (DLS) and Baishaling (BSL), while Wuzhishan (WZS), Huishan (HS), Bawangling (BWL), and Jianfengling (JFL) are clustered into group II (see Figure 2). The small subgroups clustered in group I were further divided into two smaller subgroups, indicating a relatively significant genetic relationship between each other in this larger group. The aggregation of samples in the second group is relatively chaotic, suggesting that there is less genetic relationship among the individuals in this population. From this perspective, although there is some geographical isolation between *C. hainanense* resources from different areas, there is a direct and necessary relationship between clustering based on genetic distance and their geographical origin.

**Figure 2.**
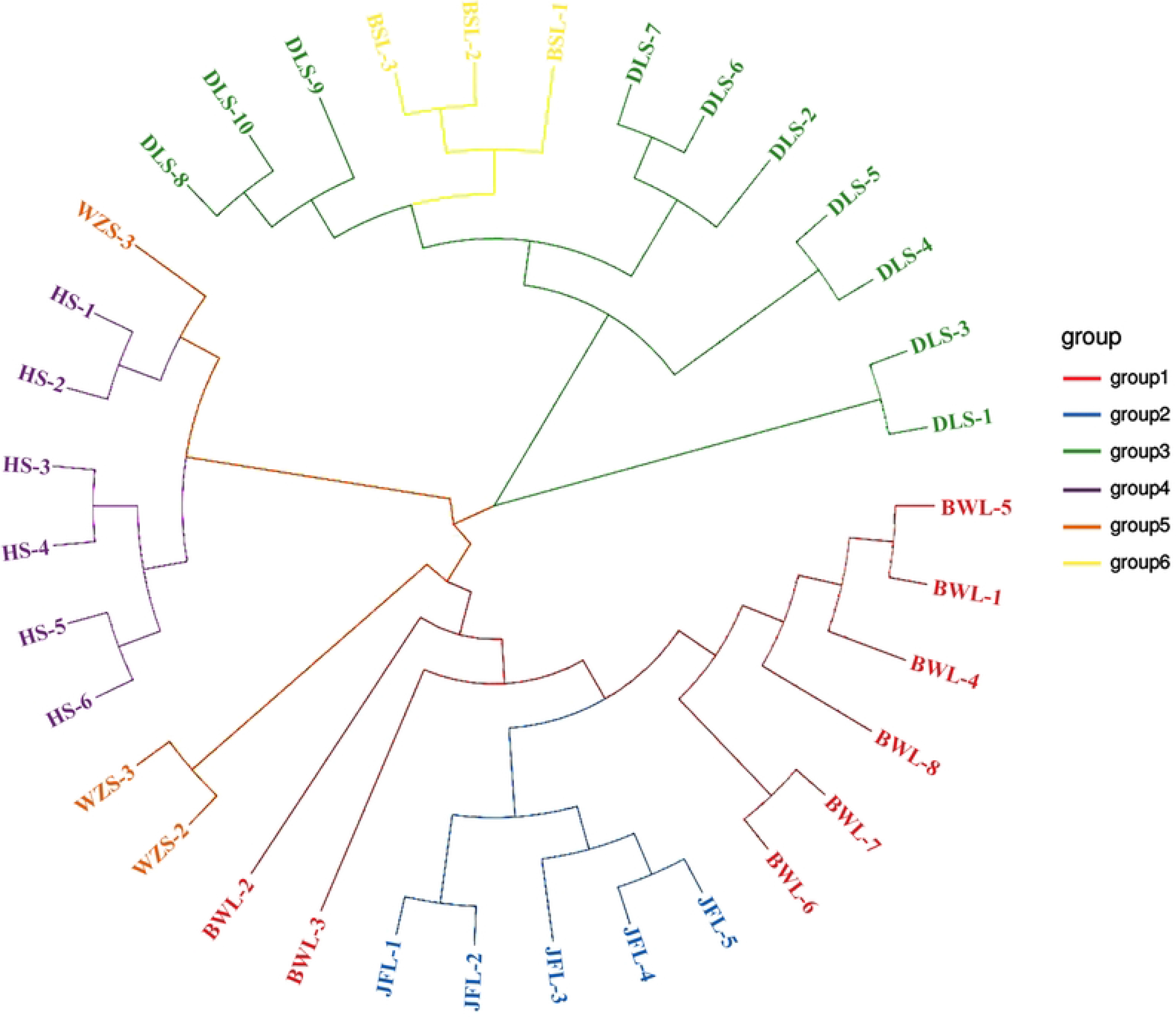
The neighbor-joining cluster of *C. hainanense* in different population

#### 3.3.2 **Analysis of population genetic structure**

When K=2, the samples from group 1 appear almost dark blue, and the samples from group 2 appear almost light purple. Upon dividing the samples into two subgroups, the samples from JFL and BWL were clustered into group 1, and the remaining samples were clustered into group 2 (Figure 3). In the cross-validation (CV) error plot (Figure 4), CV error reached its minimum value when K=1, indicating that the genetic differences among *C. hainanense* samples were relatively small, and their genetic relationships were close. Thus, it can be preliminarily inferred that the six *C. hainanense* populations in Hainan Island originated from the same ancestor and had undergone more gene exchanges. In the table of genetic differentiation coefficients among populations (Table 4), the F_st_ values among the six *C. hainanense* populations were between -0.09648 to 0.076729. There was a moderate degree of genetic differentiation (0.05 < F_st_ < 0.15) between the two populations of BSL and JFL. Furthermore, the genetic differentiation among other populations was low, indicating that differentiation was not significant (F_st_ < 0.05).

**Figure 3.**
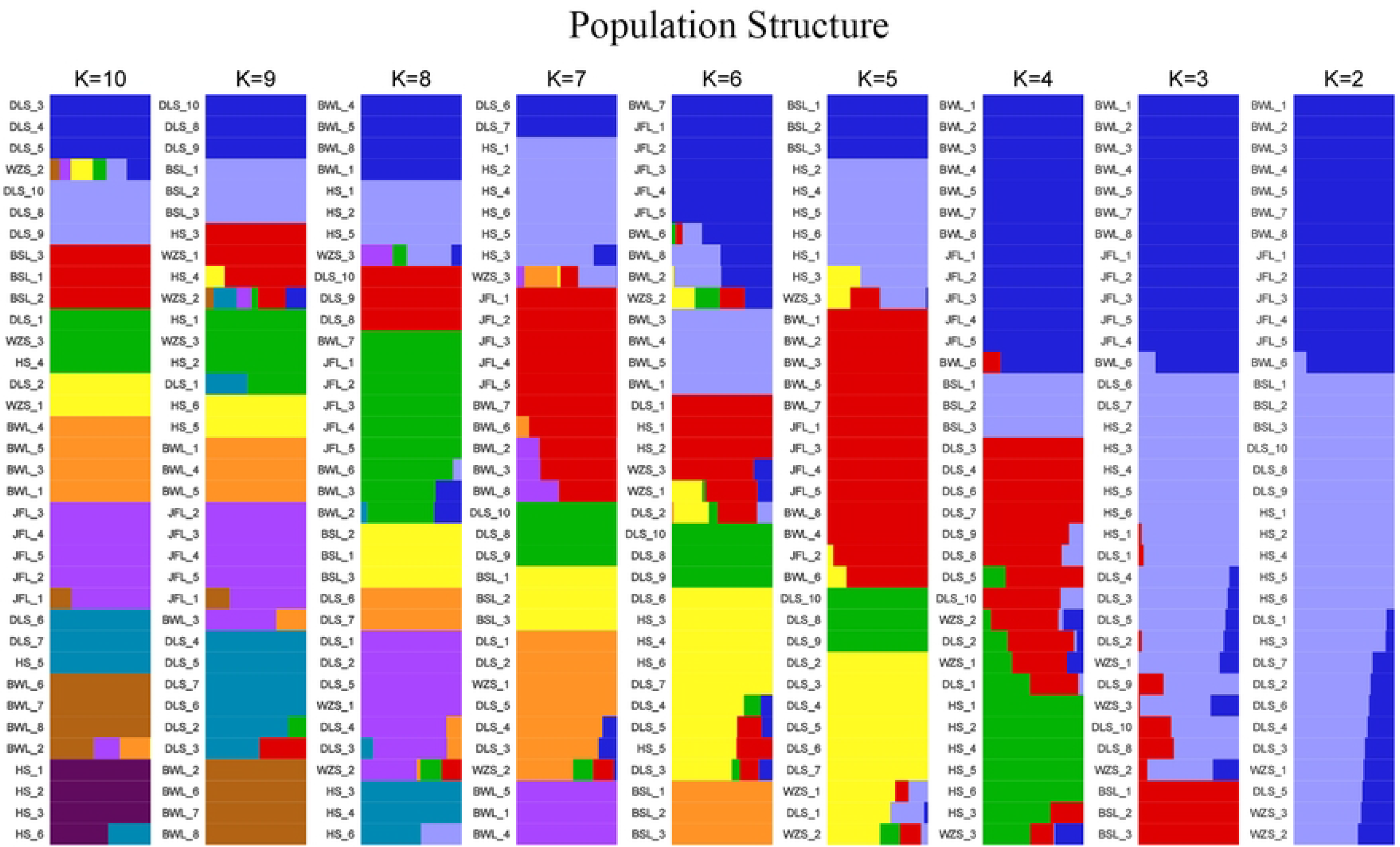
The population structure analysis on *C. hainanense*

**Figure 4.**
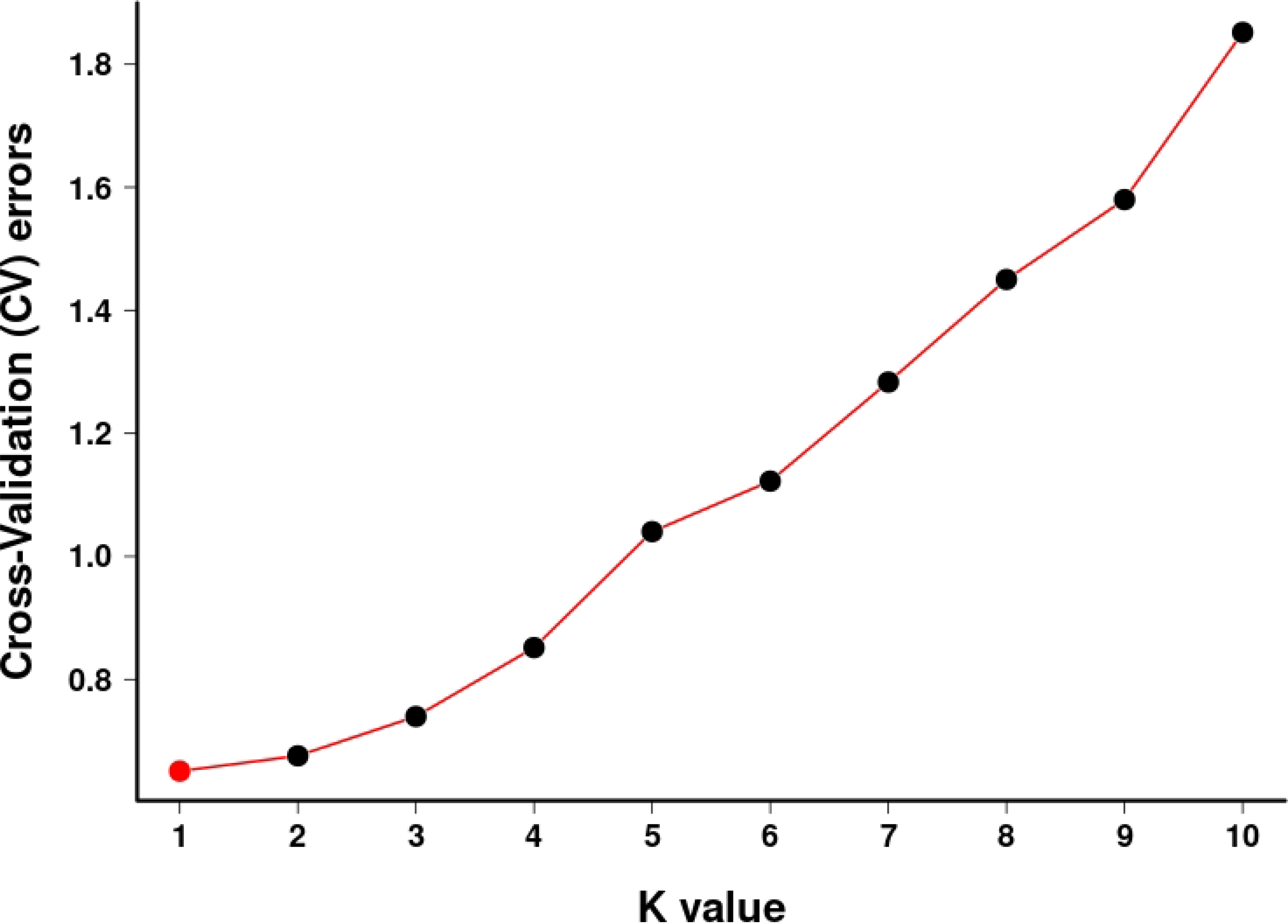
K Selection of Population Structure

**Table 4.** Genetic differentiation coefficient (F_st_: above diagonal)

|  | BWL | JFL | DLS | HS | WZS | BSL |
| --- | --- | --- | --- | --- | --- | --- |
| BWL |  | -0.06459 | -0.01902 | -0.00697 | -0.04823 | 0.025937 |
| JFL |  |  | -0.03474 | 0.007746 | -0.02721 | 0.076729 |
| DLS |  |  |  | -0.05677 | -0.09648 | -0.05853 |
| HS |  |  |  |  | -0.05853 | 0.013874 |
| WZS |  |  |  |  |  | 0.042864 |
| BSL |  |  |  |  |  |  |

### 3.4 **Principal Component Analysis Results**

Thirty-five clusters of *C. hainanense* were formed, which resulted in four independent clusters. Among these clusters, 13 samples from Jianfengling (JFL_1-JFL_5) and BWL (BWL_1-BWL_8) populations with similar genetic backgrounds, gathered to form cluster 1 (Figure 5). Six samples of *C. hainanense* from Huishan (HS_1-HS_6) and Wuzhishan (WZS_3) populations, with similar genetic backgrounds, clustered together to form cluster 2. Thirteen samples of *C. hainanense* from Diaoluoshan (DLS_1- DLS_10) and Baishaling (BSL_1-BSL_3) populations, with similar genetic backgrounds, clustered together to form cluster 3. The *C. hainanense* from the Wuzhishan (WZS_1-WZS_2) population was distant from the other three clusters, showing relatively long genetic distances, and so formed a separate cluster 4.

**Figure 5.**
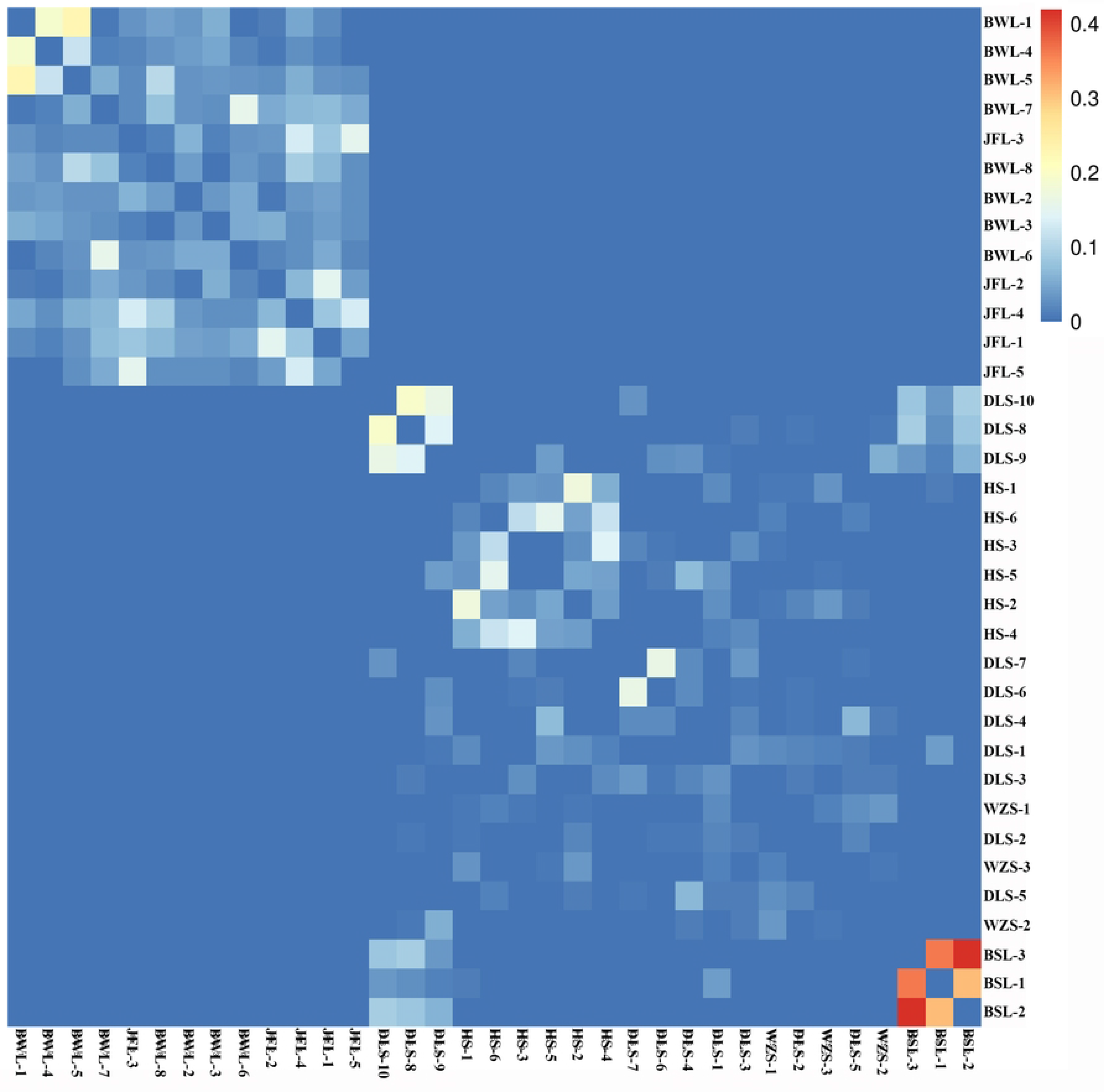
PCA analysis diagram of *C. hainanense*

### 3.5 Analysis of the genetic relationship of *C. hainanense*

We calculated the relatedness between pairs of all samples based on the SNPs (Figure 6). Kinship analysis reveals the genetic distance between samples, aiding evolutionary analysis. In the kinship heatmap, redder colors indicate closer kinship between individuals on the horizontal and vertical axes. In contrast, a red heatmap among multiple individuals suggests they may belong to a closely related family group. Conversely, bluer colors indicate more distant kinship between individuals. In the correlation heat map, the correlation coefficients of BSL_1 and BSL_2 were more outstanding than 0.4, indicating that the two samples of Bawangling were very closely related to each other. The correlation coefficients of Bawangling (BWL_1 and BWL_4), Hangluo Mountain (DLS_6 and DLS_9; DLS_7 and DLS_9; DLS_8 and DLS_10), and Huishan (HS_1-HS_6) are just between 0.2 and 0.3, indicating that some genetic exchange still exists between clusters in the case of geographical isolation (Figure 6).

**Figure 6.**
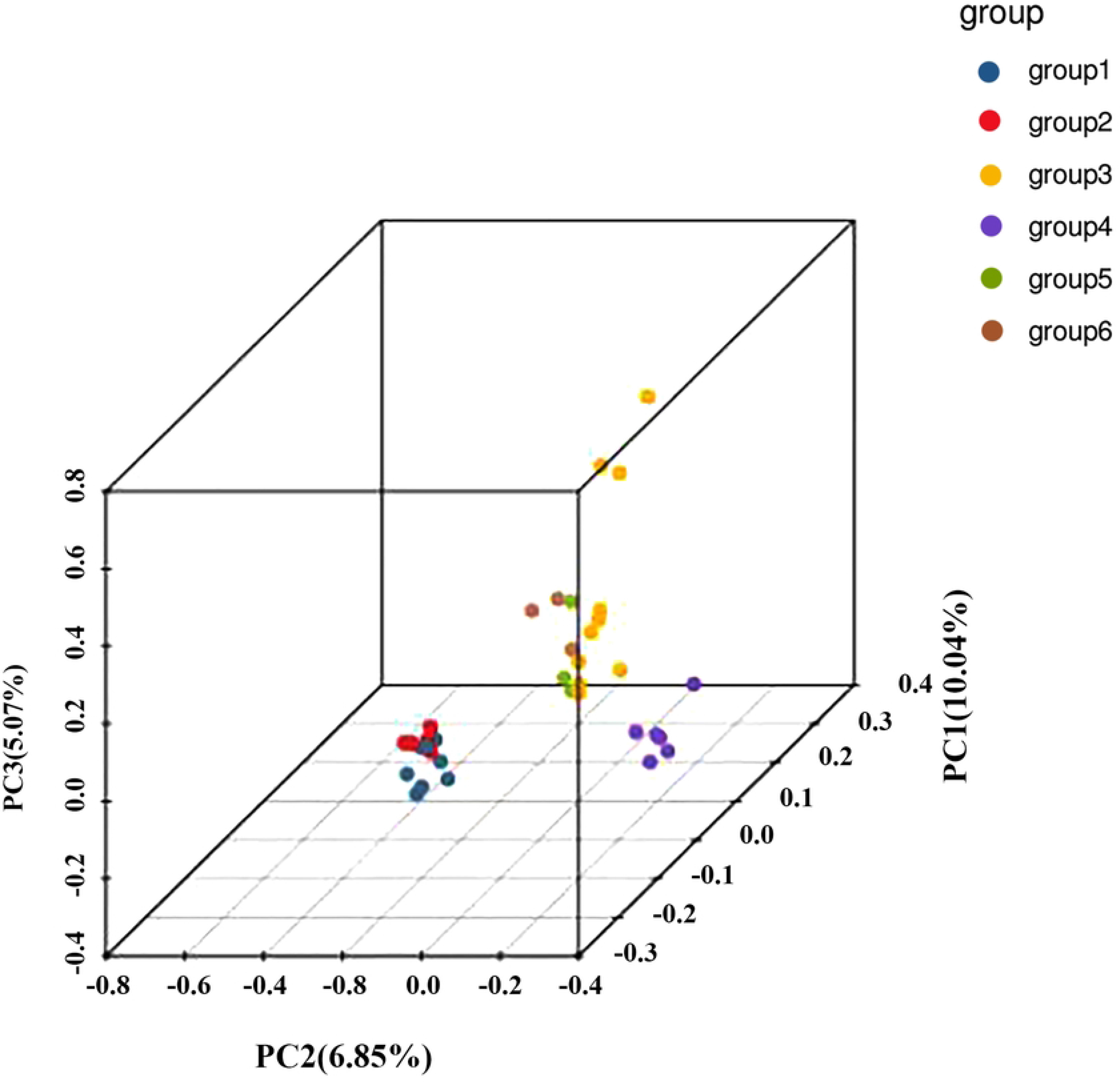
Relatedness of the Bawangling population (samples BWL_1-BWL_8), the Jianfengling population (JFL_1-JFL_5), the Luoshan population (DLS_1-DLS_10), Huishan population (HS_1-HS_6), Wuzhishan population (WZS_1-WZS_3), Baishaling population (BSL_1-BSL_3).

## 4 **Discussion**

### 4.1 Genetic diversity in *C. hainanense*

The genetic diversity of plants is usually influenced by their range, longevity, reproductive systems, seed dispersal mechanisms, and evolutionary history (Zhang et al., 2021; Nybom et al., 2004). SNP variants are the most significant and extensive type of sequence variation in plant genomes and can be easily identified by sequence alignment (Fang et al., 2014). Our study yielded 477588 high-quality SNPs through screening and filtering. *C. hainanense* has a wide ecological population in the natural wild state, but the seeds require high germination conditions in the natural environment, limiting the population. The natural wild *C. hainanense* is distributed as individuals in fragments in the natural tropical forest. So far, no clusters of communities have been found, so the population density of wild *C. hainanense* is very low, resulting in the population’s weak stress resistance and reproductive ability. Population development is long, and natural recovery is slow.

The genetic diversity of *C. hainanense* was low. The genetic diversity of *C. hainanense* was low. The finding that the genetic diversity of *C. hainanense* is low has significant implications for the conservation and management of this species. *C. hainanense* is a plant species native to Hainan Island in China. The low genetic diversity, as indicated by the heterozygosity values ranging from 10.79% to 14.55% with an average of 13.15%, suggests that this species may be at risk for several reasons. Low genetic diversity can reduce a species’ ability to adapt to environmental changes, such as shifts in climate or the emergence of new diseases or pests. A more genetically diverse population has a greater chance of possessing the necessary genetic traits to survive and adapt to new challenges. With low genetic diversity, there is an increased likelihood of inbreeding, which can lead to inbreeding depression. This is a reduction in fitness and overall health caused by the increased expression of harmful recessive genes that become more prevalent in a closely related population. Low genetic diversity can make a species more vulnerable to extinction due to its reduced adaptability and potential for inbreeding depression. When a species faces a new threat, it may not have the genetic variation necessary to evolve and survive. Genetic diversity is essential for maintaining the long-term stability and resilience of ecosystems. A species with low genetic diversity might have a reduced ability to fulfill its ecological role, which can negatively impact the overall health and functioning of the ecosystem.

WPESP in nature can lead to an increase in inbreeding due to genetic drift or bottleneck effects, which can lead to a decrease in population genetic diversity, as well as a decrease in population fitness, leaving populations unable to adapt to changing environments and facing extinction (Theodorou, et al., 2015; Nayak, et al., 2010). Xu (2022) reviewed the research on the genetic variation of 120 extremely small populations of wild plants and found that only 10 of the 44 species studied had low genetic diversity at the species level. They are *Abies ziyuanensis, thaya argyrophylla,*

*Firmiana danxiaensis, Glyptostrobus pensili, Nyssa yunnanensis, Oreocharis mileensis, Pinus squamata, Sinojackia huangmeiensis, Taxus contorta, Vatica guangxiensis* (Xu et al., 2022). At present, the distribution area of *C. hainanense* is very narrow, and the population collapse, dividing narrowly distributed may be less heritable than widely dividing plant species due to inbreeding and genetic drift (Setoguchi, et al., 2011; Zhang, et al., 2021). Low levels of genetic diversity may impair the population’s ability to adapt to new soil and photoperiodic environments during migration, thereby inhibiting the population’s adaptive potential (Yang et al., 2018). In addition, our field surveys showed that wild resources of *C. hainanense* were affected by human activities, such as logging, which led to the fragmentation of its habitat. Habitat fragmentation not only reduces the number of plant populations but also increases the spatial isolation between populations, directly or indirectly affecting species dispersal, gene exchange between populations, and species interactions, and hinders the maintenance of population genetic diversity (Aguilar et al., 2008). The findings revealed that the distribution of *C. hainanense* populations is fragmented. Many native populations have been lost due to fragmentation. Most of the *C. hainanense* populations were found in protected areas. It also shows human intervention in nature has extended into protected areas. There is an urgent need to prevent the degradation of the remaining high-quality forest ecosystems by protecting and restoring intact forests where possible, as well as by paying attention to those forest landscapes embedded in human modifications, such as those near logging fronts and those near population centers.

### 4.2 Genetic differentiation and genetic structure in *C. hainanense*

PCA analysis showed that PC1, PC2, and PC3 contributed 10.04%, 6.85%, and 5.07%, respectively. The principal component analysis’s clustering results may differ from those of the other group analyses. In PCA, the Wuzhishan population is genetically distant from other populations and forms 1 cluster separately. In addition, the cluster structure of principal component analysis, K-value selection of cluster structure, and phylogenetic tree analysis of all samples resulted in the same clusters. The fact that PCA showed the Wuzhishan population as genetically distant from other populations, forming a separate cluster, suggests that there may be unique genetic features in this population that were captured by the PCA. This could indicate that the Wuzhishan population has experienced different evolutionary pressures, gene flow patterns, or demographic histories compared to the other populations in the study. It is also important to note that the PCA clustering results may differ from other group analyses. This is because PCA is focused on capturing the maximum amount of variance in the data, while other methods, such as phylogenetic tree analysis or population structure analysis using Bayesian clustering approaches (e.g., STRUCTURE), might focus on different aspects of the data. These other methods may consider more specific evolutionary models or genetic relationships, leading to different interpretations of the population structure. Despite the differences in clustering results between PCA and other group analyses, the study found that the cluster structure of the PCA, K-value selection of the cluster structure, and phylogenetic tree analysis all resulted in the same clusters. This consistency across different analytical methods provides strong evidence for the observed population structure, lending credibility to the findings. The PCA results highlight the genetic distinctiveness of the Wuzhishan population, which may have important implications for the conservation and management of *C. hainanense*. Understanding the factors contributing to the genetic differentiation between populations can inform targeted conservation efforts aimed at preserving the unique genetic diversity found in each population. Further research on the specific genetic differences between the Wuzhishan population and others, as well as the potential environmental or ecological factors driving these differences, would provide valuable insights into the species’ evolutionary history and inform future conservation strategies.

All supported the division of 6 clusters into two clusters, so it was reasonable to divide 35 *C. hainanense* samples from 6 clusters into two clusters of 1 cluster (Diaoyu Mountain and Baisha Ridge) and 2 clusters (Wuzhishan, Huishan, Bawang Ridge, and Tsimshatsui Ridge). This division into two primary clusters could be a result of various factors, such as historical gene flow, geographical isolation, or habitat fragmentation. The two clusters may have experienced different evolutionary pressures, environmental conditions, or demographic histories that have shaped their genetic makeup. The consistency of this clustering pattern across multiple analytical methods (PCA, K-value selection, and phylogenetic tree analysis) strengthens the evidence for the observed population structure. This consistent finding implies that the division of the six populations into two primary clusters is a robust and meaningful representation of the genetic relationships among these *C. hainanense* samples.

Differences in population genetic structure are an important expression of genetic diversity. A species’ evolutionary potential and ability to withstand adverse environments depends not only on the level of genetic variation within the species but also on the genetic structure of the species (McCauley et al., 2014; Hamrick et al., 2011; Qu et al., 2004). Our results indicate that the six populations on Hainan Island are divided into two broad taxa, consistent with the principal component analysis results. A population’s genetic differentiation index (Fst) is an important parameter to measure the degree of genetic differentiation among populations. It can explain the factors that affect the genetic differentiation of populations (Zhou et al., 2022). The genetic differentiation coefficient among the populations showed that the Fst values among the six *C. hainanense* populations were between -0.09648-0.076729. There is a moderate degree of genetic differentiation (0.05 < Fst < 0.15) between the two populations of Baishaling (BSL) and Jianfengling (JFL). Furthermore, the genetic differentiation among other populations is low, and the differentiation is not noticeable (Fst < 0.05). Therefore, the genetic differentiation among *C. hainanense* populations is weak. The existing gene flow may originate from the genetic exchange between their common ancestor populations and be brought into other populations by other factors such as human factors, animal carrying, or geological factors. In addition, the topography may also affect gene flow in this species. The terrain of Hainan Island is high in the middle and low in the surrounding areas. It rises and descends to the periphery step by step. The cascade structure is prominent, and the terraces, hills, plains, and mountains form a ring-shaped layered landform (Chen et al., 2022).

The samples taken in this study were from Wuzhishan, Diaoluoshan, Jianfengling, and their surrounding forests. Geographically, the Jianfengling and Baishaling populations are far apart (>100km). However, the genetic distance between the two populations is relatively small (Fst = 0.076729), possibly due to the number of sampling populations of *C. hainanense* being too small. The population distribution of *C. hainanense* is relatively concentrated under the influence of habitat fragmentation, so the evidence is not sufficient. Therefore, the next step is to expand the scope of the sampling population for subsequent research and analysis. Limited sampling: As the study itself points out, the number of sampled populations of *C. hainanense* may be too small to accurately represent the full extent of genetic diversity and differentiation within the species. Additional sampling from other populations, particularly those that may be geographically intermediate between Jianfengling and Baishaling, could help provide a more comprehensive understanding of the genetic relationships among populations. The small genetic distance between the Jianfengling and Baishaling populations may be due to historical gene flow that occurred before the populations became geographically isolated. Over time, barriers such as geographical features, habitat fragmentation, or human activity may have restricted the movement of individuals between populations, but the genetic legacy of past gene flow could still be apparent in the current populations. *C. hainanense* may be capable of long-distance seed dispersal through mechanisms such as wind, water, or animal-mediated dispersal. Even if the frequency of long-distance dispersal events is low, they can still contribute to maintaining gene flow between geographically distant populations and reducing genetic differentiation. Human activities, such as the movement of seeds or plants for horticulture, agriculture, or reforestation efforts, may have inadvertently facilitated gene flow between the Jianfengling and Baishaling populations. This could result in the observed small genetic distance despite the large geographical separation between the populations. Small or isolated populations are more susceptible to founder effects and genetic drift, which can reduce genetic diversity and lead to increased genetic differentiation between populations. However, these processes can sometimes produce counterintuitive patterns, such as closely related populations that are geographically distant, or vice versa. To better understand the factors influencing the genetic structure of *C. hainanense*, further research is needed. This could include increasing the number of sampled populations and individuals, examining historical patterns of gene flow, and investigating potential mechanisms of seed dispersal. Ultimately, a more comprehensive understanding of the species’ genetic diversity and population structure can inform conservation strategies and help ensure the long-term survival of *C. hainanense*.

### 4.3 **Conservation and Management Strategies**

The findings of this study hold significant implications for the conservation and management of critically endangered *C. hainanense* populations within fragmented habitats in Hainan. Based on the observed patterns of genetic diversity, population structure, and inbreeding, the following conservation and management strategies are recommended:

1. Habitat protection and restoration: Prioritize the preservation of existing habitats and restore degraded areas to improve habitat quality and connectivity. Enhancing connectivity between fragmented habitats will facilitate gene flow among C. hainanense populations, promoting genetic diversity and reducing the risk of inbreeding.
2. Assisted gene flow and population augmentation: Introduce individuals from genetically diverse populations into small, isolated populations to increase genetic diversity and reduce inbreeding. This approach should be undertaken cautiously, considering potential ecological and genetic risks, such as outbreeding depression and disruption of local adaptations.
3. In-situ conservation: Efforts to protect the natural habitats of *C. hainanense* should be prioritized. This includes maintaining or establishing protected areas, enforcing regulations to prevent habitat destruction, and promoting sustainable land use practices.
4. 4. Ex situ conservation: Establish ex situ conservation programs, such as seed banks and living collections in botanical gardens, to preserve the genetic diversity of C. hainanense. These efforts can serve as a genetic reservoir for potential reintroduction or population augmentation initiatives in the future.
5. 5. Monitoring and adaptive management: Implement long-term monitoring programs to track changes in genetic diversity, population structure, and habitat conditions. Utilize the collected data to inform adaptive management strategies, ensuring the conservation efforts remain effective and responsive to emerging threats or changing circumstances.
6. 6. Community engagement and education: Involve local communities in conservation efforts by raising awareness about the importance of preserving C. hainanense and its habitat. Promote sustainable land use practices and develop community-based conservation initiatives to empower local stakeholders in the protection and restoration of the species’ habitat.
7. 7. Legal protection and policy development: Strengthen the legal protection status of C. hainanense and its habitat by incorporating the species into national and regional conservation plans. Develop and enforce policies that minimize habitat destruction, such as regulating land-use change, deforestation, and infrastructure development within the species’ range.
8. 8. Collaborative research and information sharing: Foster collaboration among researchers, conservation practitioners, and policymakers to facilitate the exchange of knowledge, data, and best practices in the conservation of C. hainanense. Encourage interdisciplinary research that integrates genetics, ecology, and social science to develop comprehensive conservation strategies.
9. 9. Climate change adaptation: Consider the potential impacts of climate change on C. hainanense populations and their habitats. Develop proactive conservation measures that enhance the species’ resilience to climate change, such as assisted migration, habitat restoration in areas with suitable future climatic conditions, and incorporation of climate change projections into spatial conservation planning.

By implementing these conservation and management strategies, we can contribute to the preservation and restoration of the critically endangered *C. hainanense* and its fragmented habitats in Hainan, while also informing efforts to protect other endangered plant species facing similar challenges.

### 4.4 **Implications for conservation**

Wild plants with extremely small populations are usually highly inbred. The characteristics of low transmission diversity and high frequency of genetic drift need to be confirmed. Genetic rescue increases genetic diversity, improves fitness, and enhances adaptation to maintain the species’ long-term survival (Sun et al., 2021). Instead of increasing the number of surviving individuals by collecting inbred seeds or asexual cuttings of *C. hainanense*, researchers should focus on designing artificial hybridization strategies to reduce inbred offspring. One of the measures to protect the *C. hainanense* population is to set up multiple protection sites to protect natural populations and their surrounding habitats. The second is to strengthen the gene flow between populations, such as constructing artificial ex-situ protection populations that should be obtained from as many different populations as possible. During ex-situ conservation, the exchange of seeds and seedlings between populations should be increased to create conditions for gene exchange and recombination artificially.

### 4.5 **Limitations and Future Research Directions**

Despite the valuable insights generated from this study, certain limitations and future research directions should be acknowledged:

1. Sample size and representation: The limited number of samples (35) and populations (six distinct cohort groups) included in this study may not entirely capture the full extent of genetic diversity and population structure of C. hainanense. Future studies should aim to increase sample sizes and cover a broader range of fragmented habitats to provide a more comprehensive understanding.
2. Gene flow and landscape connectivity: This study did not investigate the gene flow and landscape connectivity among the fragmented habitats. Understanding how landscape features influence gene flow between populations can provide crucial information for developing conservation corridors and habitat restoration strategies. Future research should incorporate landscape genetic approaches to investigate the impact of habitat fragmentation on gene flow in C. hainanense populations.
3. 3. Functional genetic diversity: The assessment of genetic diversity in this study primarily focused on neutral genetic markers (SNPs). However, functional genetic diversity, which reflects the genetic variation underlying ecologically important traits, is also crucial for species survival and adaptation. Future studies should explore functional genetic diversity by incorporating candidate genes or whole-genome sequencing approaches.
4. 4. Long-term monitoring and adaptive management: The dynamics of genetic diversity and population structure can change over time due to various factors, such as climate change, anthropogenic disturbances, and random genetic drift. Long-term monitoring of C. hainanense populations is essential for detecting changes in genetic diversity and adjusting conservation strategies accordingly.

## 5. **Conclusion**

This study provides valuable insights into the genetic diversity and population structure of the critically endangered C. hainanense within fragmented habitats in Hainan. The findings contribute to our understanding of the impact of habitat fragmentation on genetic diversity in plant populations and inform conservation efforts for other endangered plant species within fragmented ecosystems.The genetic diversity of *C. hainanense* was low, and the heterozygosity of the six populations of *C. hainanense* ranged from 10.79% to 14.55%. The average heterozygosity was 13.15%. Six *C. hainanense population* into the population (Diaoluoshan, Baishaling) and population (Wuzhishan, Huishan, Bawangling, Jianfengling) the two population. In addition, the genetic differentiation among *C. hainanense* populations is weak.

## Author Contributions

Conceptualization, Y.K.C., M.M.N.; methodology, Y.K.C., H.L.Z., M.M.N.; software, L.Z., H.L.Z., Y.K.C., M.M.N., T.T.W., T.T.L and Q.Z.;validation, M.M.N.; formal analysis, L.Z., H.L.Z., Y.K.C., M.M.N., T.T.W., T.T.L and Q.Z.; investigation, Y.K.C., and L.Z.; resources, Y.K.C.; data curation, Y.K.C.; writing—original draft preparation, L.Z., H.L.Z., Y.K.C., M.M.N., T.T.W., T.T.L and Q.Z.; writing—review and editing, L.Z., H.L.Z., Y.K.C., M.M.N., T.T.W., T.T.L and Q.Z.; visualization, Y.K.C.; supervision, Y.K.C.; project administration, Y.K.C.; funding acquisition, Y.K.C. All authors have read and agreed to the published version of the manuscript. The data presented in the study are deposited in the Dryad repository.

## Data Availability Statemen

National Science Foundation of Hainan Province The data that support the findings of this study are openly available in the Science Data Bank at https://www.scidb.cn/s/6ZZNn2.

